# The geometry of the distance-decay of similarity in ecological communities

**DOI:** 10.1101/200212

**Authors:** Joshua Ladau, Jessica L. Green, Katherine S. Pollard

## Abstract

Understanding beta-diversity has strong implications for evaluating the extent of biodiversity and formulating effective conservation policy. Here, we show that the distance-decay relationship, an important measure of beta-diversity, follows a universal form which we call the piecewise quadratic model. To derive the piecewise quadratic model, we develop a new conceptual framework which is based on geometric probability and several key insights about the roles of study design (e.g., plot dimensions and spatial distributions). We fit the piecewise quadratic model to six empirical distance-decay relationships, spanning a range of taxa and spatial scales, including surveys of tropical vegetation, mammals, and amphibians. We find that the model predicts the functional form of the relationships extremely well, with coefficients of determination in excess of 0.95. Moreover, the model predicts a phase transition at distance scales where sample plots are overlapping, which we confirm empirically. Our framework and model provide a fundamental, quantitative link between distance-decay relationships and the shapes of ranges of taxa.

## 1. Introduction

Ecological diversity is typically partitioned into three components: alpha, gamma, and beta-diversity. Alpha and gamma diversity, which quantify the species richness at local and regional scales, respectively, follow well-characterized patterns: for instance, the scaling of species-area relationships. Species-area relationships characterize the observed number of species with increasing sampling area, and are one of the most frequently studied patterns in nature. While a number of different shapes have been proposed (Rosenzweig [1995], Tjorve [2003], Lomolino [2001]) it is generally accepted that the species-area relationship follows a triphasic pattern, with a rapid species gain at local scales, followed by a leveling off at intermediate scales, and finally an accelerating increase in species number with area at the very largest, continental scales (Preston [1960], Williams [1964], Brown [1995], Rosenzweig [1995], Hubbell [2001], O’Dwyer and Green [2009]). At the intermediate spatial scales, a striking pattern that has been identified across decades of empirical studies is the power-law curve introduced by Arrhenius [1921].

By contrast, beta-diversity, which gives the turnover of community composition through space, is not known to follow well-characterized patterns. This may be because beta-diversity has been studied for less time compared to other components of diversity. Whittaker [1960] introduced the earliest measure of beta-diversity over one hundred years after Watson [1847] plotted the first species-area curve to quantify alpha diversity ([reviewed in Rosenzweig, 1995, Lomolino, 2001]). In recent years, there has been a dramatic increase the number of publications concerned with beta-diversity ([reviewed in Gaston et al, 2007, Tuomisto, 2010a,b]), including the development of new theory to predict patterns of beta-diversity (Harte and Kinzig [1997], Chave and Leigh [2002], Plotkin and Muller-Landau [2002], Morlon et al [2008]), as well as empirical papers to test those predictions (Harte et al [1999], Condit et al [2002], Tuomisto et al [2003], Green et al [2004], Morlon et al [2008]).

Despite these advances, there remain few generalizations in the literature regarding beta-diversity. For instance, distance-decay relationships show a decay of similarity between pairs of communities as a function of the distance between them (Nekola and White [1999], Soininen et al [2007]). But these relationships are not universally of an obvious functional form, and they are known to depend on the index used to quantify community similarity. Without a good model for distance-decay, it is difficult to make inferences or predictions based on observed data. This lack of knowledge comes at a great cost, given the importance of beta-diversity to practical issues of conservation planning (McKnight et al [2007], Buckley and Jetz [2008]) and to conceptual issues that are relevant to understanding the magnitude and variability of biodiversity (Novotny et al [2007], Qian and Ricklefs [2007]).

In this paper, we show that beta-diversity – measured as the distance-decay relationship – follows a universal functional form. We derive this function for a large class of community similarity metrics by making parsimonious assumptions about the geographic ranges of taxa. The geographic range of a taxon, here defined as the areas over which a taxon is is found across a landscape, has long been recognized as a fundamental building block that shapes the biodiversity and biogeography of ecological communities (Gaston and Blackburn [2000], Lomolino [2001]). However a theoretical framework linking range distributions to the distance-decay relationship does not exist. Here, we show that if ranges can be approximated by polygons, then distance-decay relationships will follow a piecewise quadratic model. Our model is general, enabling estimation of distance-decay curves for a number of commonly employed community similarity measures. We then show that this model fits distance-decay data from six disparate ecological systems extremely well. Our results establish a fundamental, quantitative link between the geometry of ranges and distance-decay relationships.

## 2 Using Geographic Ranges to Model the Distance-Decay Relationship

We propose a general model for how community composition similarity decays with increasing distance. This model enables accurate estimation of beta-diversity patterns from ecological sampling data in which pairs of plots separated by a range of distances are compared in terms of the presence and absence of taxa.

Our approach to modeling distance-decay is novel in two ways. First, we devise a method to avoid certain challenges of directly modeling indices of community similarity. Typically these indices are ratios of three quantities: the numbers of taxa (i) shared between communities, (ii) exclusive to exactly one community, and (iii) occurring in at least one community (Koleff et al [2003], Tuomisto [2010a,b]). Because averages of ratios are difficult to model, we first model separately how these three quantities scale with geographic distance, and then directly compute the standard ratio-based indices of similarity. This two-step approach enables us to identify a universal relationship between similarity and distance that only emerges when the three components of similarity indices are modeled individually. The second unique aspect of our approach is that we use results from analytic geometry in order to derive the functional form of the distance-decay. This geometric interpretation of distance-decay leads us to uncover several important ways that study design can strongly influence estimates of beta-diversity.

### 2.1 Geometric Translations Underlie the Distance-Decay Relationship

In this section we introduce the notation, terminology, and concepts used in our mathematical results. Consider a pair of sampling plots positioned randomly within a region, *Y*, which contains the ranges of the taxa of interest (Figure 1). Four key ideas, presented here, help to establish our main results, which are described in the next two sections.

**Fig. 1.**
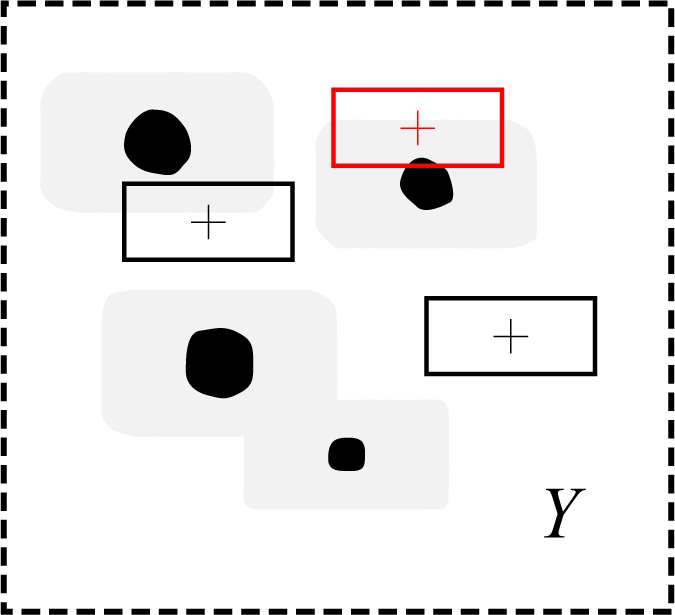
Example of a *range* and *Minkowski sum* of a range and a plot. In this example, the region *Y* (e.g., a planar region of the Pacific Ocean) is taken to be a rectangular area denoted by the dashed rectangle. *Y* is assumed to entirely contain the ranges of the taxa of interest. The black regions show a hypothetical range of a taxon; e.g., the spatial region occupied by a taxon of bacteria at 1 m beneath the ocean surface. Three possible placements of identically shaped sampling plots are given (small rectangles with plus signs at their centers). Roughly speaking, the Minkowski sum of the range and plot is the region such that the center of a sampling plot falls in the Minkowski sum if and only if the plot overlaps the range of the taxon; compare the placement of the sampling plot highlighted in red, which overlaps the the taxon range, to the other two placements which do not. The Minkowski sum depends both on the geometry of the range of the taxon, and the dimensions and orientation of the sampling plot. Our use of Minkowski sums is specified more precisely in Analytic Results and Online Resource 1.

First, for a single pair of plots separated by distance *δ*, we define an indicator variable 1_*k*_, which equals 1 if taxon *k* occurs in both of the plots, and 0 otherwise. The number of taxa shared between the plots is simply the sum *∑*_*k*_ 1_*k*_ over taxa of the indicator variables 1_*k*_. It follows that the expected number of taxa shared between pairs of plots separated by distance *δ* is the mean of these sums over many pairs of such plots. Since the mean of a sum is equivalent to the sum of the means, the expected number of shared taxa at distance *δ* is the sum over taxa of the per-taxon averages of 1_*k*_. Thus, to understand the scaling of the average number of shared taxa with distance, it is sufficient to understand how the per-taxon averages of 1_*k*_ scale with distance. Hence, we can consider one taxon at a time, which simplifies our proofs.

The second important idea we use is that because 1_*k*_ is binary (0 or 1), the average of 1_*k*_ equals the probability that 1_*k*_ equals 1; i.e., the probability that the taxon occurs in both of the plots. Thus, in order to model the relationship between shared taxa and distance, it is sufficient to understand the scaling of this probability with distance.

Our third observation is that a simple geometric rule can be established for determining when a taxon occurs in both plots. We assume that the spatial distribution of each taxon remains essentially fixed while taxa are being surveyed and that present taxa are reliably detected (though detection probabilities can easily be incorporated into the model). Hence, the taxon occurs in both of the plots if and only if both plots overlap the range of the taxon, where the “range” of the taxon is defined as the spatial region occupied by individuals of the taxon. Moreover, a plot overlaps the range of the taxon if and only if the center of the plot falls within the range of the taxon or a region adjacent to the range, which we collectively term the *Minkowski sum* of the range and plot (Figure 1 and section 6.1 of Analytic Results). Hence, the taxon occurs in both of the plots if and only if the centers of both plots fall within the Minkowski sum.

Lastly, we observe that conditional on the centers of the plots being distance *δ* apart and offset by angle *θ* (Figure 2A), the centers of both plots lie in the Minkowski sum if and only if the center of the plot 1 lies in the overlap of the Minkowski sum and the translate of the Minkowski sum by the vector (−*δ* cos *θ, −δ* sin *θ*) (Figure 2B). This overlap will hereafter be denoted 𝒪_*δ,θ,k*_. We note that in practice the designation of plot 1 (versus plot 2) is of course arbitrary. But we use the language “plot 1” and “plot 2”, because the statistical sampling framework in our proofs treats one plot as the first and conditional on its location randomly chooses a distance *δ* and angle *θ* to designate the location of the second plot. In the following section, we first derive results conditional on *δ* and *θ*, and then integrate out *θ*, providing a simple relationship between similarity and the distance *δ*.

**Fig. 2.**
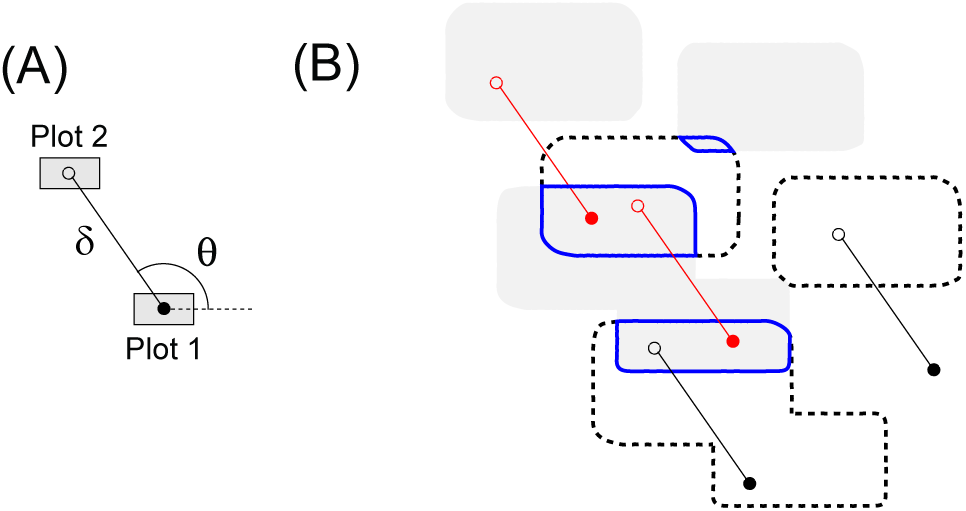
Translations and the co-occurrence of a taxon. (A) Definitions of relevant variables. Filled and open circles represent the locations of the centers of plot 1 and plot 2, respectively. The variables *δ* and *θ* refer to the distance between the plots and angle between them, respectively. (B) A condition for the taxon to occur in both plots. The gray regions represent a hypothetical Minkowski sum for the taxon (see Figure 1); the dashed regions its translate by (−*δ* cos *θ, −δ* sin *θ*). The taxon occurs in both plots if and only if the centers of both plots lie in the Minkowski sum (Figure 1). This figure shows that conditional on *δ* and *θ*, the latter condition is met if and only if the center of plot 1 occupies the intersection of the Minkowski sum and its translate, outlined in blue. Four possible locations of the plots are diagrammed; only in the cases highlighted in red do both centers lie in the Minkowski sum.

### 2.2 Taxa-Level Model

Putting these observations together, we are able to derive a specific functional form for the relationship between the expected number of shared taxa between two plots and the distance *δ* separating them. Given that the plots are distance *δ* apart and offset by *θ*, the average value of 1_*k*_ equals the probability that the center of the first plot lies in *𝒪*_*δ,θ,k*_. Clearly, *𝒪*_*δ,θ,k*_ − and therefore this probability − depends on the distance between the plots (*δ*) and their offset (*θ*). It also depends on how plots are distributed across the region *Y*, the shape and size of the plots, and the range of taxon *k*. It is necessary to make assumptions about these latter three quantities. The assumptions that our theory makes are unrestrictive. With regard to the sampling, intuitively, if plots are concentrated in atypical areas of *Y*, a different distance-decay relationship may result than if plots are randomly distributed across *Y*. Thus, our theory assumes the following.

Assumption 1: Plots are uniformly and independently distributed across the union of *Y* and, to mitigate boundary effects, a region adjacent to *Y* (defined in Sections 6.1 and 6.2 of Analytic Results). We will derive results conditional on both plots being within *Y*.

Assumption 2: Each plot is rectangular, of the same dimensions, and oriented identically. (This assumption could be generalized by following similar arguments to those presented here to develop theory for other plot configurations.)

Assumption 3: The range of taxon *k* consists of a set of polygons and points, whose Minkowski sums are at least *δ*_0_ units apart and, in the case of polygons, whose Minkowski sums are at least *δ*_0_ units “wide” (see sections 6.1 and 6.2 of Analytic Results and Online Resource 2). Here, *δ*_0_ is defined as the maximum distance over which it is of interest to predict the distance-decay relationship.

These assumptions are appropriate for many data sets: sampling is often roughly uniform, plots are distributed independently, and the ranges of many taxa can reasonably be approximated with polygons. Moreover, simulation results suggest that our theory may be robust to moderate deviations from the assumptions (data not shown). It is possible, therefore, that our theory is a special case of a more general theory requiring even weaker assumptions.

To derive predictions about distance-decay relationships, Assumption 1 implies that conditional on plot 1 being in *Y* and the plots being distance *δ* apart and offset by *θ*, the probability of the center of plot 1 falling in an arbitrary region within *Y* is proportional to the area of that region (Lemma 2 of Analytic Results and Online Resource 1). Hence, by the above observations, to show how the average of 1_*k*_ scales with distance, it is necessary to show how the area of 𝒪_*δ,θ,k*_ scales with distance.

To evaluate this scaling, for simplicity, assume here that the range of the taxon consists of a single polygon meeting Assumption 3 (the full theory given in Analytic Results and Online Resource 1 allows for more general ranges). Under this assumption, for *δ* less than an upper bound and given *θ*, the area of 𝒪_*δ,θ,k*_ is a quadratic polynomial (i.e., polynomial of degree 2 or less) in *δ* (Lemma 9 of Analytic Results and Online Resource 1).Thus, the average of 1_*k*_ conditional on the plots being distance *δ* apart and offset by angle *θ* is

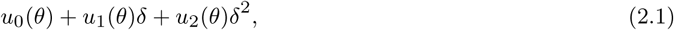

where *u*_0_, *u*_1_, and *u*_2_ are functions that depend on the angle *θ* between plot 1 and plot 2, but not on the distance *δ* separating the plots. Therefore, we see that the relationship between expected number of shared taxa and distance is a quadratic function in distance, in which the coefficients do not depend on distance.

This result suggests that the distance-decay relationship for a single taxon may be easily estimated, especially if we can also remove the dependence of the coefficients in Equation (2.1) on the angle between plots. In order to obtain the relationship between community similarity and distance alone, we derive an equation that integrates the angle *θ* out of Equation (2.1). This integration shows that conditional on the plots being distance *δ* apart, the average value of 1_*k*_ is indeed a quadratic polynomial in *δ* in which the coefficients do not depend on *δ* or *θ* (Lemma 13 of Supplementary File 1). This result provides a simple quadratic function relating the probability that two plots both contain a given taxon (i.e., the average of 1_*k*_ for taxon *k*) to the distance between the plots.

### 2.3 Community-Level Model

Our taxa-level model can be used to derive a community-level model for the expected number of shared taxa between plots as a function of distance. If the range of each taxon consists of a polygon, we have shown that the expected (i.e., average over plots) value of 1_*k*_ is a quadratic function in distance *δ* for each taxon *k*. As described above, the fact that an average of sums is equivalent to a sum of averages means that we can sum these taxa-level equations over taxa to obtain an expression for the expected value of *Σ*_*k*_ 1_*k*_, which is the expected number of shared taxa. And since each taxa-level equation is a quadratic function of *δ* with coefficients that do not depend on *δ*, the coefficients in these models are simply added together to produce a single quadratic equation in *δ*:

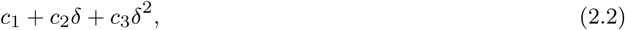

where *c*_1_ to *c*_3_ do not depend on *δ*.

If taxa have more complicated ranges, consisting of multiple polygons and/or isolated points, it is necessary to add a non-quadratic term *c*_4_*v*(*δ*) to the polynomial, where *c*_4_ is a constant and *v*(*δ*) is a function with no unknown parameters that depends on the dimensions of the sampling plot (Section 6.1 and Theorem 1 of Analytic Results). The correction term *v*(*δ*) is zero for distances larger than the diagonal distance between two opposite corners of a sampling plot, and it takes on non-zero values at smaller distances that depend on the relationship between *δ* and the lengths of the two sides of the plot. The addition of a non-zero *v*(*δ*) term at small distances *δ* results in a biphasic distance-decay relationship that takes a different form at small compared to larger distances. Inclusion of this correction term enables modeling of distance-decay at dimensions smaller than the sampling plot (i.e., where sample plots overlap). This range has not typically been explored but might be important, for example, in studies of microbial diversity where sampling areas may be practically constrained to be much larger than relevant micro-environments.

Finally, these results are conditional on plot 1 being in *Y*. To condition on both plots falling in *Y*, two approaches are possible, one exact and one approximate. In the exact approach, the shape of *Y* needs to be known. When this is the case, it is necessary to correct for boundary effects by dividing by another function, *v*(*δ, Y*), which has no unknown parameters and depends on the shape of *Y* (Section 6.1 of Analytic Results). This function *v*(*δ, Y*) adjusts for the fact that after sampling plot 1 in *Y*, we sometimes cannot sample all possible plots distance *δ* from plot 1 (i.e., plots with centers at all *θ ∈* [0, 2*π*]), because some of these will be outside *Y*. The correction depends on *δ*, since plots that are close together are less likely to hit the boundary of *Y* than plots that are further apart. Importantly, implementing this correction requires that taxa occur in roughly the same density at all distances from the boundary of *Y*. The details of this approach are given in Theorem 2 of Analytic Results.

When the shape of *Y* is unknown, an approximate approach is available. It can be shown that when *δ* is small relative to the dimensions of *Y*, the boundary correction described above becomes negligible (Theorem 3 of Analytic Results). Thus, no knowledge of the shape of *Y* is required because the function *v*(*δ, Y*) can be omitted. Moreover, homogeneity of taxon densities across *Y* does not need to be assumed. For many data sets – e.g., surveys from tropical forest census plots – it may be difficult to know *Y*, which is defined as a region entirely containing the ranges of the taxa, and *δ* is likely small relative to the dimensions of the ranges of taxa. Thus, we focus on the approximate approach in the following, bearing in mind that the exact approach is available when *Y* is known.

Incorporating these correction terms into Equation (2.2), we find that the average number of taxa occurring in both plots is the piecewise quadratic law

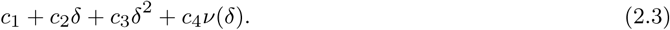

Thus, we have derived a simple community-level model for the relationship between the average number of taxa shared between plots and distance. This model is a quadratic polynomial with the correction term *v*(*δ*) to account for taxon ranges that are not single polygons. This model is appropriate at distance-scales that are small relative to the dimensions of *Y* (Theorem 3 of Analytic Results). Because of the *v*(*δ*) term,the resulting distance-decay model is approximately piecewise quadratic, i.e., the combination of different quadratic functions over different distance rages. This piecewise function enables accurate modeling of biphasic distance-decay relationships (See Section 4).

So far, we have focused only on the average number of shared taxa. Similar mathematics can be used to show that the average number of taxa occurring in exactly one plot and the average number of taxa occurring in at least one plot follow related piecewise quadratic functions (Equations (6.19) and (6.20) in Analytic Results). Details are given in Theorem 3 of Supplementary File 1. Then, all three results can be combined to derive distance-decay relationships for a wide range of different community similarity metrics, since these metrics are typically simple functions of the numbers of shared taxa, taxa in exactly one plot, and taxa in at least one plot. Here we demonstrate this approach with four commonly used indices of similarity: the (i) Sorenson, (ii) Jaccard, (iii) Whittaker (Whittaker [1960]), and (iv) Wilson and Shmida (Wilson and Shmida [1984]) indices. We refer to these indices as *β*_*s*_, *β*_*j*_, *β*_*w*_, and *β*_*t*_, respectively, following Koleff et al [2003].

For example, the Sorenson index *βs* is defined as twice the number of taxa shared between plots divided by the sum of the number of taxa in the plots. According to our theory, the numerator of this expression is given by twice Equation (6.18). The denominator is given by the sum of Equation (6.18) and Equation (6.17) (Theorem 3 of Analytic Results). Therefore, the average of *βs* can be approximated by the ratio of the functions,

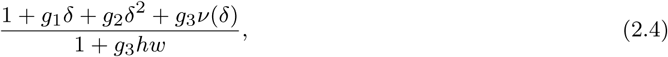

where *h* and *w* denote the (known) height and width of the sampling plots, respectively and *g*_*i*_ is defined as *c*_*i*+1_*/c*_1_ for *i* = 1, 2, 3. Likewise, the average of the Jaccard index *β*_*j*_ is approximated by

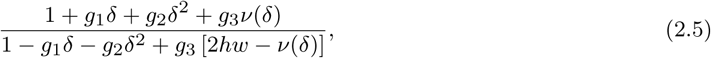

and the averages of *β*_*w*_ and *β*_*t*_ are approximated by

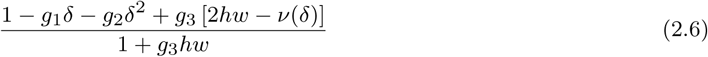

and

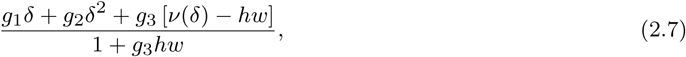

respectively.

We conclude this section by noting several observations of our approach. First, our estimator of each index is the ratio of the expected value of its numerator and the expected value of its denominator. For example, we estimate the expected value of the Sorenson index as the ratio of twice the expected number of shared taxa divided by the expected number of taxa in both plots. This is not exactly the same as the expected value of the index, because ratios of expectations are not equivalent to expectations of ratios. There are two reasons that we take this approach. First, the expectation values of the indices are not always well defined. This problem follows from the fact that plots have nonzero probability of being devoid of taxa. Hence, the denominator of an index can be zero for a given pair of plots, leading to an undefined value of the ratio and therefore an undefined value of the expectation. The second motivation for our approach is that using the ratio of expectations is common practice in the beta-diversity literature ([e.g., Morlon et al, 2008]).

Second, it is interesting that equations (2.4) to (2.7) are of similar form: the ratio of *v*(*δ*) and polynomials of degree two or less. Since *v*(*δ*) takes on different functional forms for different ranges of distances *δ* (see above), the distance-decay model for each of the similarity indices is approximately a piecewise quadratic function in distance *δ*.

Finally, we note that the equations for estimating different similarity indices share parameters. For instance, *g*_1_ is the same in Equations (2.4) and (2.5). These shared parameters reflect the correlation between the quantities that we model (i.e., the union is the sum of the intersection and the symmetric difference). In total, the model has three parameters: *g*_1_, *g*_2_, and *g*_3_. We estimate *c*_1_ to *c*_4_ once and then use these to estimate *g*_1_ to *g*_3_. Since these shared estimators are plugged into the equations for different indices, different measures of similarity computed on the same data set will be appropriately correlated with one another.

## 3 Methods

To test our model, we employed several survey data sets where the true distance-decay relationship can be computed directly. Fore each data set, we used data *in silico* resampling to simulate the process of conducting a beta-diversity study in which pairs of plots at varying distances from one another are sampled and the species present in each plot are recorded. Using the simulated data, we fit our model and compared model predictions to the true distance-decay relationship.

### 3.1 Data Sets

We analyzed six published data sets: three surveys of neotropical rainforest vegetation (Barro Colorado Island, Cocoli, and Sherman; Condit [1998], Hubbell et al [1999, 2005]), a survey of vegetation in a 16*×*16 meter region of serpentine grassland in California (Green et al [2003]), and digitized atlases of mammals and amphibians in a 29 × 22 degree region of South America (IUCN et al [2008], Patterson et al [2007]). These data sets vary greatly in taxonomic scope and spatial scale (256 m^2^ to over 10^6^ km^2^). Each data set gives the spatial distribution of species within the respective taxon in the specified region.

### 3.2 Simulated Sampling

For each data set, we constructed distance-decay relationships by simulating the uniform and independent placement of pairs of square sample plots separated by a wide range of different distances *δ* and recording the species occurring in each plot (see Supplementary File 4). The dimensions of the plots were 25 *×* 25 m for the Barro Colorado Island vegetation data, 10 *×* 10 m for both the Cocoli and Sherman vegetation data, 1.5 *×* 1.5 m for the serpentine vegetation data, and 2 *×* 2 degrees for both the amphibian and mammal data. Analysis with different heights yielded qualitatively similar results (not reported here). For each data set we considered 100 values of the distance *δ*. For each value of *δ*, we iterated the algorithm 750 times, using 250 observations to independently model each of the three piecewise quadratic laws (Equations (6.18), (6.19) and (6.20)).

### 3.3 Model-Fitting

Relative to size of the ranges of most of the taxa, the dimensions of the surveys that we considered were small. For example, the dimensions of Barro Colorado Island survey are 500 × 1000 m, but the ranges of most plants found therein are greater than 100 km^2^. Thus, the short-distance model discussed above was appropriate, under the assumption that the distance-decay curves found for the surveys were representative of those that would have been found by sampling across the entire ranges (at short distances).

Model fitting was performed using a weighted least squares procedure (Supplementary File 4). The response function for the indices of similarity in Equations (2.4) to (2.7) are nonlinear. Rather than fit these response functions directly, we fit the response functions given in Equations (6.18) to (6.20) (Analytic Results), which are linear, to obtain estimates 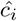 of *c*_*i*_, *i* =1,…4. We then used 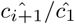 as estimates of the three parameters *g*_*i*_, *i* = 1, 2, 3 (recall that *g*_*i*_ is defined as *c*_*i*+1_*/c*_1_). Although the former estimates may not have minimized the nonlinear error sum of squares for Equations (2.4) to (2.7), very high coefficients of variation indicated that they were nearly minimized.

### 3.4 Goodness of Fit

Because the data sets are censuses in which we know the true overlap between species in all pairs of plots, we can compare the predictions of the proposed model to the observed distance-decay relationship. To do so, we used two methods for quantifying goodness of fit. The first method was purely descriptive: a coefficient *WLS* of determination. Because our regression values were weighted, we used 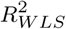 values (Willett and Singer [1988]). Typical weighted coefficients of variation would yield similar general conclusions, although their use is less appropriate here because of the repeated measures structure in our sampling framework (i.e., many pairs of plots at each distance). The second method with which we quantified goodness of fit was a lack of fit test. To perform these tests, we bootstrapped the observed data (Stute et al [1998]).

A complication in evaluating goodness of fit is that our theoretical results predict how the ratio of average values scales with distance, rather than how the average of ratios scales. As noted in Section 2.3, we are therefore modeling the different similarity indices by taking ratios of two functions with shared parameters. Hence, to evaluate goodness of fit we need to consider these two functions in each ratio separately. To do so, we first compute the sample mean of the numerator and denominator of an index across all plots at a given distance *δ*. Then, we use the ratios of these sample means across different values of *δ* to evaluate goodness of fit. To estimate standard errors for the ratios, we used the delta method.

## 4 Results

### 4.1 Analysis of Beta Diversity in Different Ecosystems

To investigate how well our piecewise-quadratic law describes distance-decay data, we consider six published survey data sets (Methods). For each data set, we derived the true distance decay relationship and compared this to the predicted relationship from our model. To derive the model prediction for each data set, we simulated a distance-decay study by distributing *in silico* pairs of plots separated separated by a range of different distances (Methods). Using the simulated data, we estimated the three model parameters *g*_*i*_, *i* = 1, 2, 3 and derived curves relating each of the four community similarity metrics described above (Figure 3 and Supplementary File 3) to distance. We then examine the fit of these models to the true distance-decay curve.

**Fig. 3.**
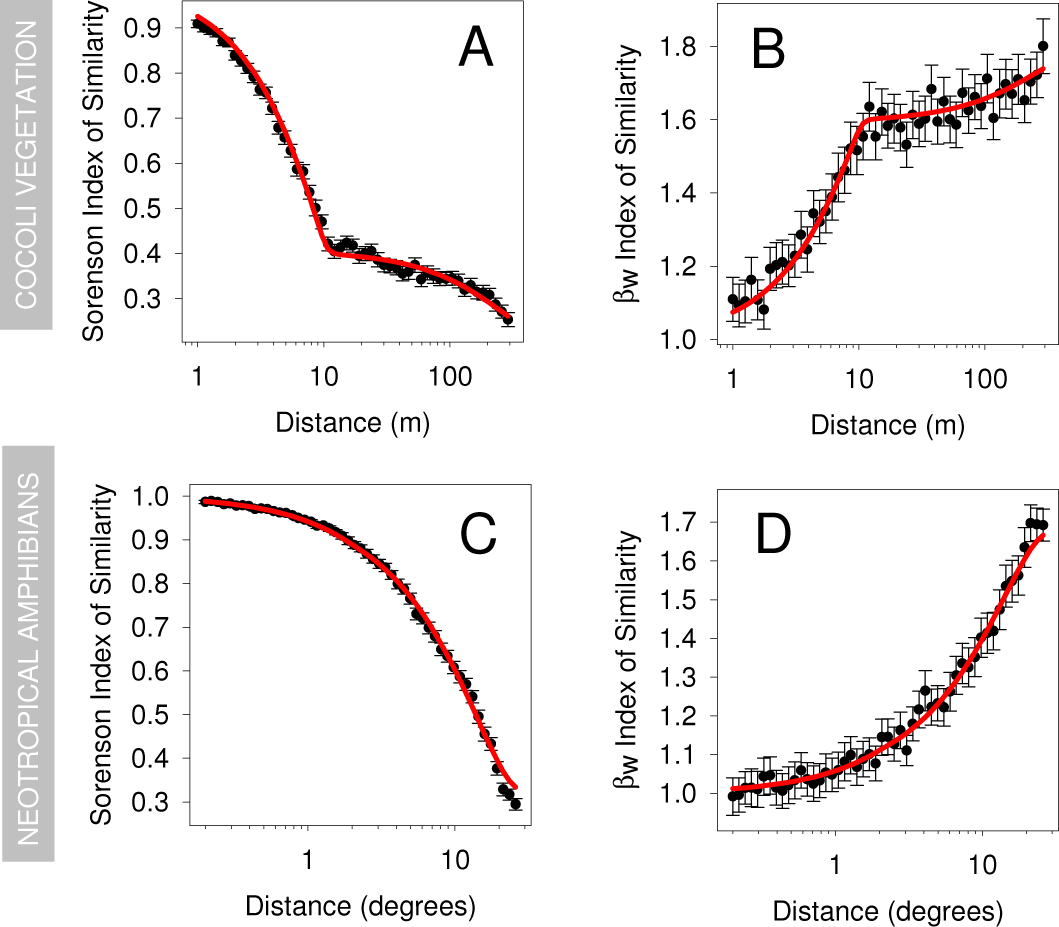
Assessing the distance-decay model via computer simulation using survey data. Two indices of similarity, *β*_*s*_ and *β*_*w*_ are compared on the Cocoli Vegetation (A, B) and neotropical amphibian data (C, D). In each graph, the independent variable is the distance between the center of pairs of censused plots, while the dependent variable is the indicated index of similarity. The black dots and error bars (2 standard errors) give the true values of the distance-decay relationship across simulated pairs of plots. The red lines give the model fits. The coefficients of determination (*R*^2^) for the fits in all graphs are at least 0.97. These results show that we can accurately estimate distance-decay for very different similarity indices (panels A vs. B and C vs. D) across ecosystems with very different distance-decay patterns (panels A and B vs. C and D).

### 4.2 Model Assessments

We find that the piecewise quadratic law fits the data sets exceptionally well for each of the four community similarity metrics. Predictions of the Sorenson, Jaccard, and Wilson-Shmida indices of similarity account for more than 97% of the observed variation. Predictions for the Whittaker index account for more than 93% of the variation. Although a lack of fit analysis shows significant deviations, these deviations are relatively minor and due primarily to large sample sizes and very high statistical power (Table 1 and Figure 3). Because our model has three parameters, it is reasonable to ask whether such strong agreement between true and predicted distance-decay relationships is remarkable. To address this question, we simulated additional data sets from a variety of parametric models that are not derived from ecological data. This sensitivity analysis shows that with the sample sizes we analyzed, poor fits can easily occur (Supplementary File 3). Thus, we conclude that the close fit of the observed data from different ecosystems to the proposed piecewise-quadratic law indicates that this model is indeed an appropriate description of the underlying beta-diversity.

**Table 1.** Goodness of fit results. Coefficients of determination for six data sets and four measures of beta-diversity are shown. For each data set, a model with three parameters was fit once. This model predicts the scaling of all four indices of diversity. *β*_*s*_, *β*_*j*_, *β*_*w*_, and *β*_*t*_ refer to the Sorenson, Jaccard, Whittaker (1960), and Wilson and Shmida (1984) indices of similarity, respectively. Except for the neotropical amphibians data set, lack of fit tests were significant, likely reflecting slight deviations from the model and high statistical power.

| Data Set | Response Variable |  |  |  |
| --- | --- | --- | --- | --- |
| | $\beta_s$ | $\beta_j$ | $\beta_w$ | $\beta_t$ |
| BCI Vegetation | 0.991 | 0.989 | 0.978 | 0.991 |
| Cocoli Vegetation | 0.996 | 0.993 | 0.973 | 0.996 |
| Sherman Vegetation | 0.994 | 0.994 | 0.979 | 0.994 |
| Neotropical Amphibians | 0.999 | 0.993 | 0.980 | 0.999 |
| Neotropical Mammals | 0.997 | 0.996 | 0.988 | 0.997 |
| Serpentine Vegetation | 0.992 | 0.978 | 0.935 | 0.992 |

### 4.3 Comparison of Distance-Decay Patterns

Our model enabled accurate estimation of a wide variety of distance-decay patterns (Figure 3 and Figures 3 to 7 in Supplementary File 3). Overall, the data sets analyzed showed the expected decay of community similarity with distance, but the shape of this decay differed in notable ways between ecosystems. The slopes of the distance-decay relationships were much greater for the vegetation surveys than the neotropical mammal and amphibian distributions. Although variability in dispersal and life history traits could account for these differences, an equally likely hypothesis is that they result from differences in the resolution of the data. The resolution of the location data for the vegetation is on the order of 1 cm, while for the mammals and amphibians, it is on the order of 100 km. In the former case, if a species was known to occur within a roughly 1 cm grid cell, then that species was recorded as being present therein, while in the latter case, grid cells were roughly 100 km. With data being pooled at large resolutions, it is not surprising that similarity decays slowly. However, within the vegetation data sets, where data resolution was constant, similarity decayed more quickly with distance in the temperate ecosystem than the tropical one, an observation that accords with previous studies.

We also found that our model accounts well for observed phase shifts in the distance-decay patterns of these diverse ecosystems. In all of the vegetation data sets, we observed a pronounced phase shift at distances equal to the height of the sampling plot, as predicted by our theory. Our fitted model was able to accurately capture this biphasic pattern. We also found a slight distance-decay phase shift in the mammal and amphibian data. But high variance in the data prevented this prediction from being confirmed or accurately compared to the shifts observed in the vegetation data sets.

## 5. Discussion

We have shown that the distance-decay relationship, a common measure of beta-diversity, can be modeled by a piecewise quadratic function that is easily and accurately estimated from sampling plot data. Until now, a universal form for distance-decay was not known. In fact, the variety of distance-decay patterns observed across different ecosystems suggested that an underlying beta-diversity relationship, like that of the species-area relationship for alpha diversity, might not exist. However, our theoretical results provide some insight into why such a relationship may indeed exist. The distance-decay model that we derive is essentially quadratic. But it deviates from a simple quadratic function in an important way.

Our theory predicts that distance-decay relationships are piecewise quadratic functions. That is, up to a distance determined by the dimensions of the sampling plot, they follow one quadratic polynomial, while at larger distances, they follow a different quadratic polynomial. The two phases result from effects that isolated individuals have on the probability that a taxon will be observed in two plots separated by distance *δ*. These effects only contribute to the distance-decay relationship when plots overlap. We account for this contribution of isolated individuals at small scales with a model term *v*(*δ*) that takes on different forms depending on *δ* and the dimensions of the plots. This biphasic pattern that emerges from our proofs is evident in the data. We found that distance-decay relationships estimated from simulating sampling across a range of ecosystems show abrupt changes in slope at the distances predicted by our theory. These estimated phase changes mirror those seen in the true distance-decay relationships for these same data sets. Moreover, additional simulations using different sized sampling plots resulted in the distances of the phase change shifting as we would predict (data not shown). That our theory correctly predicts a heretofore unseen phase transition at these distances is strong evidence of its validity.

Examining distance-decay relationships at distance scales smaller than the dimensions of sampling plots is not typically done [e.g., Morlon et al, 2008]. However, new insights can no doubt be gained by considering beta-diversity patterns at these small scales, particularly for small organisms where restricting distance-decay studies to non-overlapping plots may exclude very relevant distance ranges. Consider, for example, studies of soil microbes. The minimum size of a sampling plot is typically fixed – for instance, the diameter of soil cores may be constrained by the sampling apparatus and physical limitations. But, spatial scales smaller than a single soil core are likely relevant to microbial ecology. By sub-sampling from cores to generate data from overlapping “plots” and applying our biphasic piecewise quadratic model, distance-decay patterns at the relevant, small scales could be accurately estimated, as illustrated for Cocoli vegetation in panels A & B of Figure 3 where distances less than 10m represent scales smaller than the plot width.

Accounting for plot dimensions is critical to accurately estimating distance-decay at small scales, but this modeling subtlety was not known previously. It was revealed by our approach, which treats taxon ranges as fixed (at least on the timescale of the study) and focuses on modeling the random distribution of sampling plots on the landscape. Rather than explicitly focusing on sampling, models of distance-decay have previously been based solely on assumptions about the distributions of taxa. For instance, the methods of (Harte and Kinzig [1997], Harte et al [1999]) assumed that species’ spatial distributions are fractal, and in the Poisson cluster process proposed by Morlon et al [2008], individuals are modeled as dispersing in a specific stochastic manner. Our theory breaks from this approach. We assume that the distributions of individuals meet certain general criteria. But, the random stochastic quantities in our framework are the locations of the sampling plots – quantities that so far have not been explicitly modeled in theories predicting distance-decay relationships. The validity of the predictions tested here suggest that random sampling over generalized ranges underlies distance-decay relationships observed in disparate ecological systems.

We focused on model predictions for cases when distances between plots are small relative to the size of the ranges of taxa. In doing so, we assumed that the distance decay curves constructed from each data set were representative of those that would have been constructed by sampling the entirety of *Y* (at short distances). However, at distances close to the dimensions of the regions surveyed for each data set, plots can only be placed close to the boundary of the surveyed region. For instance, in an extreme case, if the surveyed region is a rectangle and the distance between plots is the length of the diagonal of the region, then the plots can only be placed at opposite corners of the region. Thus, because of the increased reliance on just those observations along the boundary at large distances, the similarity values observed at these distances have greater uncertainty associated with them than those at short distances (which are based on many pairs of plots). Estimating the magnitude of this uncertainty is challenging. One approach is to check whether the boundaries of the surveys differ from the centers – if they do, then pairs of observations along the boundary are suspect. For the vegetation surveys, the boundaries appear similar to the centers, but one could imagine circumstances – for instance, surveying an island whose boundary consists of a beach – wherein the boundaries would be dissimilar to the centers. In most studies, this uncertainty is not explicitly considered. Thus, our geometric approach to modeling distance-decay has highlighted an important consideration for designing a beta-diversity study, which, if ignored, can cause unknown uncertainty about how community similarity scales with distance.

We treated the extent of the ranges of taxa as unknown quantities, approximating boundary effects as being negligible (Theorem 3 of Analytic Results and Supplementary File 1). When the ranges, and thus *Y*, are known, more accurate predictions are possible. In these cases, it is necessary to include an additional model term *v*(*δ, Y*), and assume that taxa are distributed homogeneously at all distances from the boundary of *Y* (Theorem 2 of Analytic Results and Supplementary File 1). These complications arise from boundary effects analogous to those discussed above, but occurring along the boundaries of *Y* rather than the smaller surveyed region. Importantly, the extent of these boundary effects not only depends on the distance between plots, but also the shape of the region *Y* containing the taxon ranges; hence, the model term *v*(*δ, Y*). It is a new prediction that the distance-decay relationship can be affected by the shape of *Y*. In addition, another exact approach to the boundary problem is to condition on just one plot being within *Y* rather than both plots. This approach requires different distributional assumptions, and leads to a different model term that is a function of *δ* and *Y*.

Our theory makes predictions additional to those that we tested. For instance, distance-decay relationships for all subsets of taxa (e.g., just the Asteraceae in the 50 Ha BCI survey region) should follow the same law, a prediction that appears to be valid based on preliminary analyses. Additionally, our work suggests that if the surveyed region is a volume rather than a two dimensional area – for instance a cubic kilometer of ocean – then the quadratic polynomials in Equations (2.3) to (2.7) become cubic polynomials, with corresponding modifications to the function *v*(*δ*). Investigation of these predictions will lead to additional understanding of the appropriateness of our theory and universality of patterns of beta-diversity across different environments. Distance-decay relationships arise in other fields as well; for instance, in linguistics and economics. Investigation of these relationships from other fields may yield further insight into the mechanisms underlying the spatial scaling of diversity.

The excellent model fit that we observe across a range of ecosystems relies upon a few simple assumptions about the study design. Most importantly, we assume that (i) taxon ranges can be approximated by certain polygons and isolated points, and (ii) sampling plots are uniformly and independently distributed across the study area and (iii) distances between plots are small relative to the size of the ranges of taxa. Starting from these assumptions, we were able to derive a universal distance-decay model by applying a variety of geometric and algebraic principals. One of the results we use – that the area of intersection of two translated polygons locally depends on the distance between them as a quadratic function of distance – was derived in computer science literature, where it is has implications for packing and object recognition algorithms (Mount et al [1996]). That such a disparate field can be productively brought to bear on challenging ecological problems highlights the power interdisciplinary approaches to ecology, and the need for applying new mathematics to ecology.

Taxon ranges are a fundamental building block for understanding biodiversity and biogeography patterns. Ecologists have long recognized that there is enormous variation in the sizes of geographic ranges of individual taxa, and have sought to understand the causes and consequences of this variation. For example, it has been documented that oftentimes the frequency distribution of geographic ranges within an ecological community is unimodal and right skewed ([reviewed in Gaston and Blackburn, 2000, Lomolino, 2001]), and that geographic ranges increase with increasing latitude (Rapoport and Bariloche [1982]). In addition to the sometimes regular variation in the sizes of geographic ranges, regular patterns in the shapes of ranges (i.e., their perimeter-to-area ratio) among different taxon have also been documented (Brown [1995]). While substantial effort has been put forth to understand how these types of regularities, and the overlap of ranges, ultimately influence alpha-diversity, little is known about how the shapes of geographic ranges influences beta-diversity. In this paper, we began with assumptions about the shapes of the ranges of taxa, and showed that these assumptions implied the functional form of distance decay-relationships. This logic could be turned around: beginning with distance-decay relationships, attributes of the ranges of taxa could potentially be predicted. Ongoing work indicates that this is indeed the case. As such, distance-decay relationships could provide powerful tools for inferring attributes of taxon ranges. The theory developed here is a crucial step in quantitatively linking taxon ranges and distance-decay relationships.

## 6. Analytic Results

### 6.1 Definitions

In the following, capital letters will denote sets, while lower case letters will denote points. Bold letters will denote random variables and vectors. Where necessary, superscripts will be used differentiate variables. Except where otherwise noted, the variables *X, X*^1^, *X*^2^, *…*will denote arbitrary subsets of ℝ^2^, while *x, x*^1^, *x*^2^, *…* will denote arbitrary points in ℝ^2^. Subsets of sets will be denoted by subscripts; e.g., *X*_1_ *⊆ X*. Likewise, coordinates of points will be denoted by subscripts; for instance, *x ≡* (*x*_1_, *x*_2_). For any set *S*, 1_*S*_ will be the characteristic function of *S*. The symmetric difference operator will be ⊖.

Let *∂X* be the boundary of *X*. Polygons will be taken as closed sets. For any polygon *𝒫*, let *x*^1^ be *adjacent* to *x*^2^ in *𝒫* if and only if the line segment joining *x*^1^ and *x*^2^ is contained in *∂𝒫*. Let *V* (*𝒫*) be the set of vertices of *𝒫*; i.e., the set of *x ∈ ∂𝒫* such that there exist *x, x ∈ ∂𝒫* that are adjacent to *x* but not each other.

Let *X* + *X* be the Minkowski sum of *X*^1^ and *X*^2^, and correspondingly, let the sum *X* + *x* denote the translate of *X* by *x*. For any *θ ∈* ℝ, let 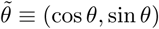. Thus, for any *δ ∈* ℝ, 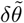 is the mapping of (*δ, θ*) from polar coordinates to Cartesian coordinates. Throughout the following, a real number *θ* will be termed an *angle* if and only if *θ ∈* [0, 2*π*]. *δ* will be used to denote distances; it will be assumed real and nonnegative. Define *d(X,*^1^ *X*^2^) as the distance between *X*^1^ and *X*^2^. Likewise, if *X* is a polygon, let d (*X*) equal the minimum distance between non-intersecting edges of *X*; otherwise define it as 0. Fixing *w* and *h* as positive numbers, let *R* be the rectangle given by

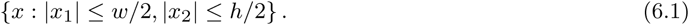

Fixing *δ*_0_ as a positive number, define a *polygon-singleton cover* as follows:

#### Definition 1.

*A set X is a polygon-singleton cover of X if and only if*

1. *X consists of polygons and singleton subsets of* ℝ^2^.
2. 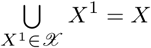.
3. 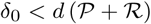 *for all polygons* 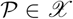.
4. 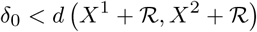 *for all distinct* 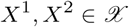.

Let *μ* be the Lebesgue measure, and *Y* a closed Lebesgue measurable subset of ℝ^2^. *Y* represents a region containing the ranges of the species of interest – for instance, the Pacific Ocean or Amazon Basin. It will be assumed that each species occurs only within *Y* (Assumption 2a, below). Let *n* denote the number of taxa that occur in *Y*, and for *k* = 1, *…, n*, let *Yk* denote the range of species *κ*; i.e., the subset of *Y* that species *κ* occupies. Each range will be taken as a fixed, non-random quantity. For any *δ*, let *B*_*δ*_ (*x*) be the closed ball (disk) centered at *x*. For convenience, let *B*_*δ*_ ≡ *B*_*δ*_ (0). Define the sample space

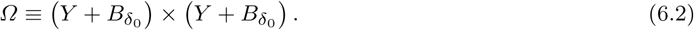

The two coordinates of each sample point represent the centers of a pair of *h × w* rectangular plots in which taxa are surveyed. Sample points will be denoted 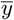, to emphasize that rather than elements of *Y* per se, they are elements of the direct product space. A sample point 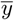 will be taken to equal (*y*^1^, *y*^2^). For *i* = 1, 2, let {*X*}^*i*^ be the event that the center of plot *i* is an element of *X*:

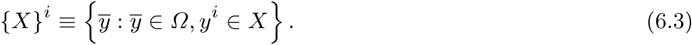

Likewise, for *k* = 1,…,*n*, let ⟨*k, i*⟩ be the event that plot *i* intersects *Y*_*k*_:

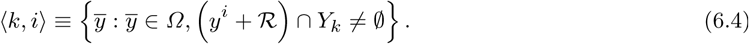

Denote complementary sets by a tick mark; e.g., ⟨*k, i*⟩^t^ is the complement of ⟨*k, i*⟩.

On *Ω*, define *T* ^*i*^ so that for 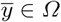

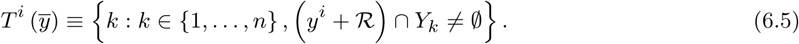

That is, 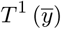 and 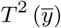 are the sets of taxa occurring in plots 1 and 2, respectively. Define the random vectors **Y**^1^, **Y**^2^ : *Ω →*ℝ^2^ so that 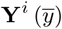 gives *y*^*i*^, *i* = 1, 2. Let 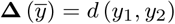. (**Δ** and *d* differ in that the domain of **Δ** is a subset of the domain of *d*.) Define arctan : ℝ^2^ *→*ℝ so that arctan (*x*) gives the signed angle between *x* and (1, 0). Let 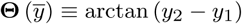.

Given a random variable or vector **V** on *Ω*, we will denote the cumulative distribution function and probability density functions of **V** by *F***_V_** and *f***_V_**, respectively. Define the real-valued function *v* so that for any *δ ∈* ℝ and closed Lebesgue measurable set *X* ⊆ ℝ^2^,

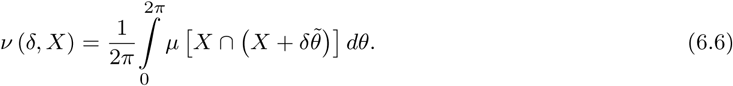

For simplicity, in the foregoing non-rigorous text, we refer to *v*(*δ, R*) simply as *v*(*δ*).

A *quadratic polynomial* will be defined as a polynomial of degree 2 or less.

### 6.2 Assumptions

Our results require two classes of assumptions:

1. [Sampling] The sampling model, *ℳ*, consists of the set of probability measures on *Ω* such that for any *P ∈ℳ*,
  a. **(a)Y**^1^ and **Y**^2^ are uniform on 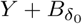.
  b. **(b)Y**^1^ and **Y**^2^ are independent.
2. [Geometry of occupied regions] For *k ∈* {1, *…, n*},
  a. *(a)Yk* + ℛ *⊆ Y*.
  b. *(b)Yk* has an polygon-singleton cover.

The set of all *n*-tuples (*Y*_1_,…, *Y*_*n*_) satisfying assumptions 2a and 2b will be denoted *Y*. In the following, *Y*_1_, …, *Y*_*k*_ will be assumed to satisfy assumptions 2a and 2b.

### 6.3 Main Results

This section presents our main results. Proofs of these results and supporting lemmas can be found in Supplementary File 1.

The first result shows that two conditional distributions are uniform. This result is key for linking the distribution of plots to the geometry of the ranges of taxa.

#### Lemma 2.

*For any δ ≤ δ*_0_, *angle θ, y ∈ Y, and P ∈ℳ, conditional on* {*Y* }1:

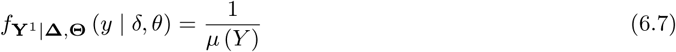

*and*

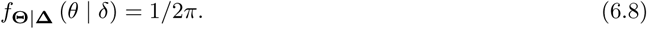

The second result gives the set of critical placements sensu Mount et al [1996]. A important attribute of this set is that it is finite.

#### Lemma 8.

*For any polygon P and δ < d* (*P*),

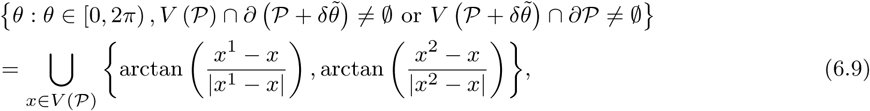

*where x*^1^ *and x*^2^ *denote the vertices adjacent to x.*

Lemma 9 shows that the expected area of an intersection is a quadratic polynomial. This result can be derived from Lemma 8, and along with Lemma 2, is used to show the piecewise quadratic model.

#### Lemma 9.

*For any polygon P,*

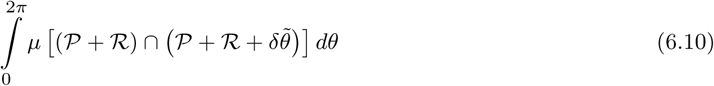

*is a quadratic polynomial in δ on* (0, *d* (*P* + *R*)), *the coefficients of which are functions of P.*

Lemma 14 quantifies edge effects. Specifically, all results are initially derived conditioning on just {*Y* }^1^. By multiplying by the factor given in Lemma 14, these results can be modified to allow conditioning on {*Y* ^*i*^}^1^ *∩* {*Y ′*}^2^.

#### Lemma 14.

*For any P ∈ M, δ < δ*_0_, *and discrete random variable* **V** *on Ω, if for all nonzero i ∈*ℕ {**V** = *i*} *∩*{*Y ′*}= *ϕ*, *then*

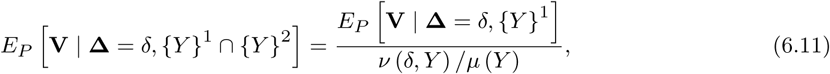

*where v is defined in* *Equation* (6.6).

For positive *δ*, the piecewise quadratic model includes the function *v*(*δ, ℛ*). Theorem 1 gives a closed form expression for this function.

#### Theorem 1.

*For any positive δ*

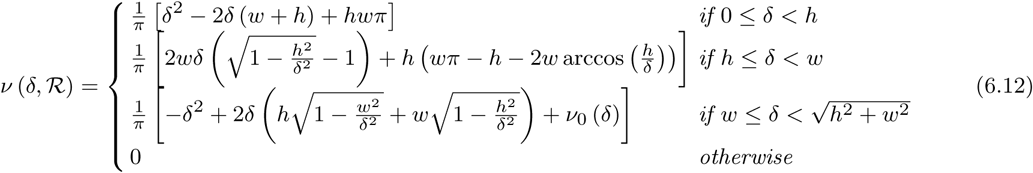

*where*

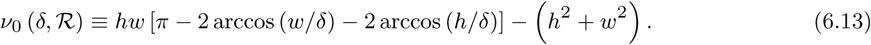

Theorem 2 gives the exact form of the piecewise quadratic model. It requires that *Y* is known, and that *δ < δ*_0_.

#### Theorem 2.

*If for all δ < δ*_0_ *and i* = 1, 2,

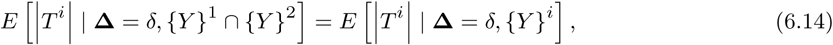

*then there exist real-valued functions c*_1_, *…, c*_4_ *on Y such that for all δ < δ*_0_ *and P ∈ M,*

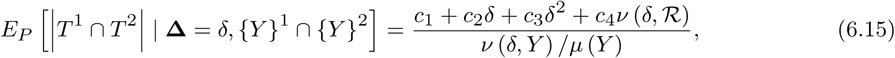

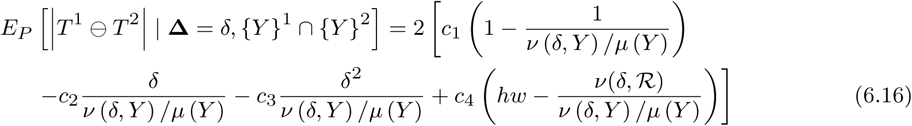

*and*

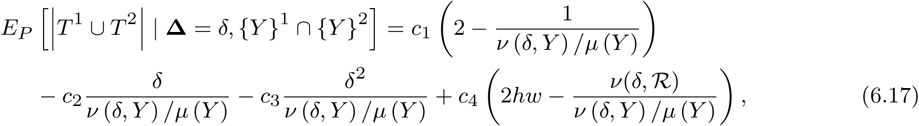

*where *v* is defined in Equation* (6.6).

Theorem 3 gives the approximate form of the piecewise quadratic model. For the approximation, *Y* does not need to be known, and substantially weaker conditions need to hold (compare the hypotheses of Theorem 3 to those of Theorem 2).

#### Theorem 3.

*In the limit μ*(*Y*)*/μ Y* + *Bδ*0 *→* 1, *there exist real-valued functions c*_1_, *…, c*_4_ *on Y such that for all δ < δ*_0_ *and P ∈ ℳ,*

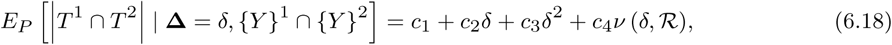

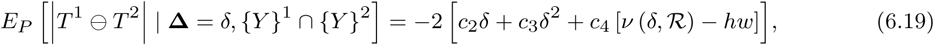

*and*

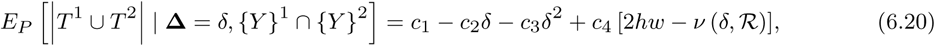

*where *v* is defined in Equation (6.6)*.

## Acknowledgements

This work was supported by the Betty and Gordon Moore Foundation (Grant 1660). Helpful suggestions and advice were kindly provided by Tony Capra, Steven Kembel, Dennis Kostka, James O’Dwyer, Samantha Riesenfeld, and Thomas Sharpton.

## References

Arrhenius O (1921) Species and area. Journal of Ecology 9:95–99

Brown J (1995) Macroecology. University of Chicago Press

Buckley L, Jetz W (2008) Linking global turnover of species and environments. Proceedings of the National Academy of Sciences 105:17,83617,841

Chave J, Leigh E (2002) A spatially explicit neutral model of beta-diversity in tropical forests. Theoretical Population Biology 62:153–168

Condit R (1998) Tropical Forest Census Plots. Springer-Verlag and R. G. Landes Company

Condit R, Pitman N, Leigh Jr E, Chave J, Terborgh J, Foster R, Nunez P, Aguilar S, Valencia R, Villa G, Muller-Landau H, Losos E, Hubbell S (2002) Beta-diversity in tropical forest trees. Science 295:666–669

Gaston K, Blackburn T (2000) Pattern and process in macroecology. Wiley-Blackwell

Gaston K, Evans K, Lennon J (2007) Scaling Biodiversity, Cambridge University Press, chap The scaling of spatial turnover: pruning the thicket, pp 181–222

Green J, Harte J, Ostling A (2003) Species richness, endemism and abundance patterns: tests of two fractal models in a serpentine grassland. Ecology Letters 6:919–928

Green J, Holmes A, Westoby M, Oliver I, Briscoe D, Dangerfield M, Gillings M, Beattie A (2004) Spatial scaling of microbial eukaryote diversity. Nature 432:747750

Harte J, Kinzig A (1997) On the implications of species-area relationships for endemism, spatial turnover, and food web patterns. Oikos 80:417–427

Harte J, McCarthy S, Taylor K, Kinzig A, Fischer M (1999) Estimating species-area relationships from plot to landscape scale using species spatial-turnover data. Oikos 46:45–54

Hubbell S (2001) The Unified Neutral Theory of Biodiversity and Biogeography. Princeton University Press

Hubbell S, Foster R, O’brien S, Harms K, Condit R, Wechsler B, Wright S, De Lao S (1999) Light gap disturbances, recruitment limitation, and tree diversity in a neotropical forest. Science 283:554–557

Hubbell S, Condit R, Foster R (2005) Barro colorado forest census plot data. URL http://ctfs.si/edu/datasets/bci

IUCN, International C, NatureServe (2008) An analysis of amphibians on the 2008 iucn red list. URL http://www.iucnredlist.org/initiatives/amphibians

Koleff P, Gaston K, Lennon J (2003) Measuring beta diversity for presence-absence data. Journal of Animal Ecology 72:367–382

Lomolino M (2001) The species-area relationship: new challenges for an old pattern. Progress in Physical Geography 25:1–21

McKnight M, White P, McDonald R, Lamoreux J, Sechrest W, Ridgely R, Stuart S (2007) Putting beta diversity on the map: Broad-scale congruence and coincidence in the extremes. PLoS Biology 10:2424–2432

Morlon H, Chuyong G, Condit R, Hubbell S, Kenfack D, Thomas D, Valencia R, Green J (2008) A general framework for the distance-decay of similarity in ecological communities. Ecology Lett 11:1–14

Mount D, Silverman R, Wu A (1996) On the area of overlap of translated polygons. Computer Vision and Image Understanding 64:53–61

Nekola J, White P (1999) The distance decay of similarity in biogeography and ecology. Journal of 26:867–878

Novotny V, Miller S, Hulcr J, Drew R, Basset Y, Janda M, Setliff G, Darrow K, Stewart A, Auga J, Isua B,Molem K, Manumbor M, Tamtiai E, Mogia M, Weiblen G (2007) Low beta diversity of herbivorous insects in tropical forests. Nature 448:692–695

O’Dwyer J, Green J (2009) Field theory for biogeography: a spatially explicit model for predicting patterns of biodiversity. Ecology Letters 12:1–9

Patterson BD, Ceballos G, Sechrest W, Tognelli M, Brooks T, Luna L, Ortega P, Salazar I, Young B (2007) Digital distribution maps of the mammals of the western hemisphere, version 3.0

Plotkin J, Muller-Landau H (2002) Sampling the species composition of a landscape. Ecology 83:3344–3356

Preston F (1960) Time and space and the variation of species. Ecology 41:612–627

Qian H, Ricklefs R (2007) A latitudinal gradient in large-scale beta diversity for vascular plants in north america. Ecology Letters 10:737–744

Rapoport E, Bariloche F (1982) Areography: geographical strategies of species. Pergamon Press

Rosenzweig M (1995) Species Diversity in Space and Time. Cambridge University Press

Soininen J, McDonald R, Hillebrand H (2007) The distance decay of similarity in ecological communities. Ecography 30:3–12

Stute W, Manteiga W, Quindimil M (1998) Bootstrap approximations in model checks for regression. Journal of the American Statistical Association 93:141–149

Tjorve E (2003) Shapes and functions of speciesarea curves: a review of possible models. Journal of Biogeography 30:827–835

Tuomisto H (2010a) A diversity of beta diversities: straightening up a concept gone awry. part 1. defining beta-diversity as a function of alpha and gamma diversity. Ecography 33:2–22

Tuomisto H (2010b) A diversity of beta diversities: straightening up a concept gone awry. part 2. quantifying beta-diversity and related phenomena. Ecography 33:23–45

Tuomisto H, Ruokolainen K, Yli Halla M (2003) Dispersal, environment, and floristic variation of western amazonian forests. Science 299:241244

Watson H (1847) Cybele Britannica: or british plants and their geographical relations. Longman and Company

Whittaker R (1960) Vegetation of the siskiyou mountains, oregon and california. Ecological Monographs 30:279–338

Willett J, Singer J (1988) Another cautionary note about r2: Its use in weighted least squares regression analysis. American Statistician 42:236–238

Williams CB (1964) Patterns in the Balance of Nature. Academic Press

Wilson M, Shmida A (1984) Measuring beta-diversity with presence-absence data. Journal of Ecology 72:1055– 1064

